# Two subsets of circulating Ly6C^lo^ monocytes distinguished by CD138 (syndecan-1) expression and Nr4a1 dependence in pristane-treated mice

**DOI:** 10.1101/2022.01.09.475578

**Authors:** Shuhong Han, Haoyang Zhuang, Rawad Daniel Arja, Westley H. Reeves

## Abstract

Chronic peritoneal inflammation following pristane injection induces lupus with diffuse alveolar hemorrhage (DAH) and pulmonary capillaritis in C57BL/6 mice. The pathogenesis involves pristane-induced microvascular lung injury. BALB/c mice are resistant to endothelial injury and DAH. Lung disease in C57BL/6 mice is abolished by depleting monocytes/macrophages. The objective of this study was to define the role of myeloid subsets in DAH. Hemorrhage and vasculitis were abolished in *Ccr2*-/- mice, indicating involvement of bone marrow-derived monocytes/macrophages. Along with Ly6C^hi^ monocytes, we found two subsets of circulating Ly6C^lo^ monocytes: one CD138^-^ and a novel CD138^+^ subset. *Nr4a1*-dependent patrolling Ly6C^lo^ monocytes maintain vascular integrity after endothelial injury. Circulating Ly6C^lo^CD138^+^ monocytes were associated with DAH and were absent in mice without DAH. They also were absent in *Nr4a1*-/- mice, whereas Ly6C^lo^CD138^-^ monocytes were unaffected. However, *Nr4a1*-/- mice were susceptible to pristane-induced DAH and lung vasculitis, suggesting that disease onset does not require Ly6C^lo^CD138^-^ monocytes. Peritoneal Ly6C^lo^CD138^+^ Mϕ were unchanged in *Nr4a1*-/- mice, indicating that they are not derived from Ly6C^lo^CD138^+^ monocytes. We conclude that pristane-induced lung microvascular lung injury stimulates a wave of *Nr4h1*-dependent Ly6C^lo^CD138^+^ patrolling monocytes in an ineffectual effort to maintain vascular integrity in the face of ongoing endothelial damage.

## Introduction

Monocytes are derived from bone marrow (BM) macrophage (Mϕ) and dendritic cell progenitor cells (MDPs), which develop into common monocyte progenitor cells (cMoPs) and then Ccr2^+^Ccr4^lo^Ly6C^+^ monocytes. Ccl2, a ligand for Ccr2, is essential for the egress of Ly6C^+^ monocytes from the BM in response to inflammation (1, 2), after which these cells can become Ly6C^hi^Ccr2^hi^Cx3cr1^lo^CD43^-^ “classical” monocytes or Ly6C^lo^Ccr2^lo^Cx3cr1^hi^CD43^+^ “non-classical” monocytes (3). CD62L, a marker expressed by newly emigrated BM-derived Ly6C^+^ monocytes, is lost with maturation (4, 5).

Pristane or mineral oil (MO) injection causes sterile peritoneal inflammation and an influx of Ly6C^hi^ monocytes, which become Ly6C^hi^ inflammatory peritoneal Mϕ (6). The chronic inflammatory response evolves into lupus after several months (7). In C57BL/6 (B6) mice, but not other strains, the onset of lupus is heralded by lung microvascular injury culminating in diffuse alveolar hemorrhage (DAH) (8) (H Zhuang, Submitted).

In contrast to the striking predominance of peritoneal Ly6C^hi^ Mϕ in pristane-treated mice, Ly6C^hi^ Mϕ are replaced by Ly6C^lo^ Mϕ in MO-treated mice and lupus does not ensue. A subset of the Ly6C^lo^ peritoneal Mϕ expresses CD138 (syndecan-1) (9). Ly6C^lo^CD138^+^ Mϕ from MO-treated mice have an anti-inflammatory phenotype and are highly phagocytic for dead cells. The significance of CD138 expression by Mϕ is unclear and it is not known whether it is expressed by other myeloid cells, such as monocytes.

Non-classical (Ly6C^lo^) monocyte development requires the transcription factor nuclear receptor subfamily 4 group A member 1 (Nr4a1) and Notch signaling (2, 10). Although both Nr4a1-independent Ly6C^hi^ monocytes and Nr4a1-dependent Ly6C^lo^ monocytes are derived from Ly6C^+^ precursors, they have different functions. In contrast to the role of Ly6C^hi^ monocytes as mediators of inflammation, Nr4a1-dependent monocytes patrol the vascular endothelium and promote the TLR7-dependent removal of damaged cells (5, 11). However, they also can migrate across the endothelium into inflamed tissues (12, 13).

We examined myeloid cell CD138 expression in the blood and inflamed peritoneum. Ly6C^lo^CD138^+^ and Ly6C^lo^CD138^-^ Mϕ both were Nr4a1-independent and Ccr2-dependent, suggesting that they are derived from Ly6C^hi^ monocytes recruited to the peritoneum. Unexpectedly, circulating Ly6C^lo^ monocytes also consisted of two subpopulations: CD138^+^ and CD138^-^. Only the Ly6C^lo^CD138^+^ monocyte subset was Nr4a1-dependent. Compared with the Ly6C^lo^CD138^-^ and Ly6C^hi^ subsets, Ly6C^lo^CD138^+^ monocytes expressed higher levels of <u>T</u>riggering <u>R</u>eceptor <u>E</u>xpressed on <u>M</u>yeloid <u>C</u>ells <u>L</u>ike <u>4</u> (Treml4), an innate immune receptor that amplifies TLR7 signaling and binds to late apoptotic and necrotic cells (14–17). Treml4 expression was regulated by Nr4a1, consistent with a role in the Ly6C^lo^ monocyte-mediated, TLR7-dependent, removal of damaged endothelial cells. Circulating Nr4a1-dependent Ly6C^lo^CD138^+^ monocytes increased substantially after pristane, but not MO, treatment, and were more abundant in B6 vs. BALB/c mice, suggesting that they may represent an ultimately ineffectual effort by the organism to repair pristane-induced chronic endothelial injury.

## Results

Ly6C^hi^ monocyte recruitment to the peritoneum following pristane or MO injection is Ccr2-dependent (6). In the peritoneum of pristane-treated mice, Ly6C^hi^ Mϕ are more abundant than CD138^+^ Mϕ, whereas CD138^+^ Mϕ are more abundant in MO-treated mice. We examined CD138 expression on circulating and peritoneal myeloid cells from mice treated with pristane or MO.

### Circulating CD138^+^ monocytes

Ly6C^hi^ monocytes are recruited to the inflamed peritoneum and become Ly6C^hi^ Mϕ, which can downregulate Ly6C and develop into Ly6C^lo^ Mϕ (6). A subset of Ly6C^lo^ Mϕ expresses CD138 (9). We examined whether Ly6C^lo^CD138^+^ Mϕ are derived from circulating CD138^+^ monocyte precursors. CD138^+^CD11b^+^ cells were found in the blood of both pristane and MO-treated mice at 14-d, but were much more abundant in pristane-treated mice **(Fig. 1)**. In contrast, peritoneal CD138^+^CD11b^+^ Mϕ were more abundant in MO-treated mice than pristane-treated mice.

**Figure 1.**
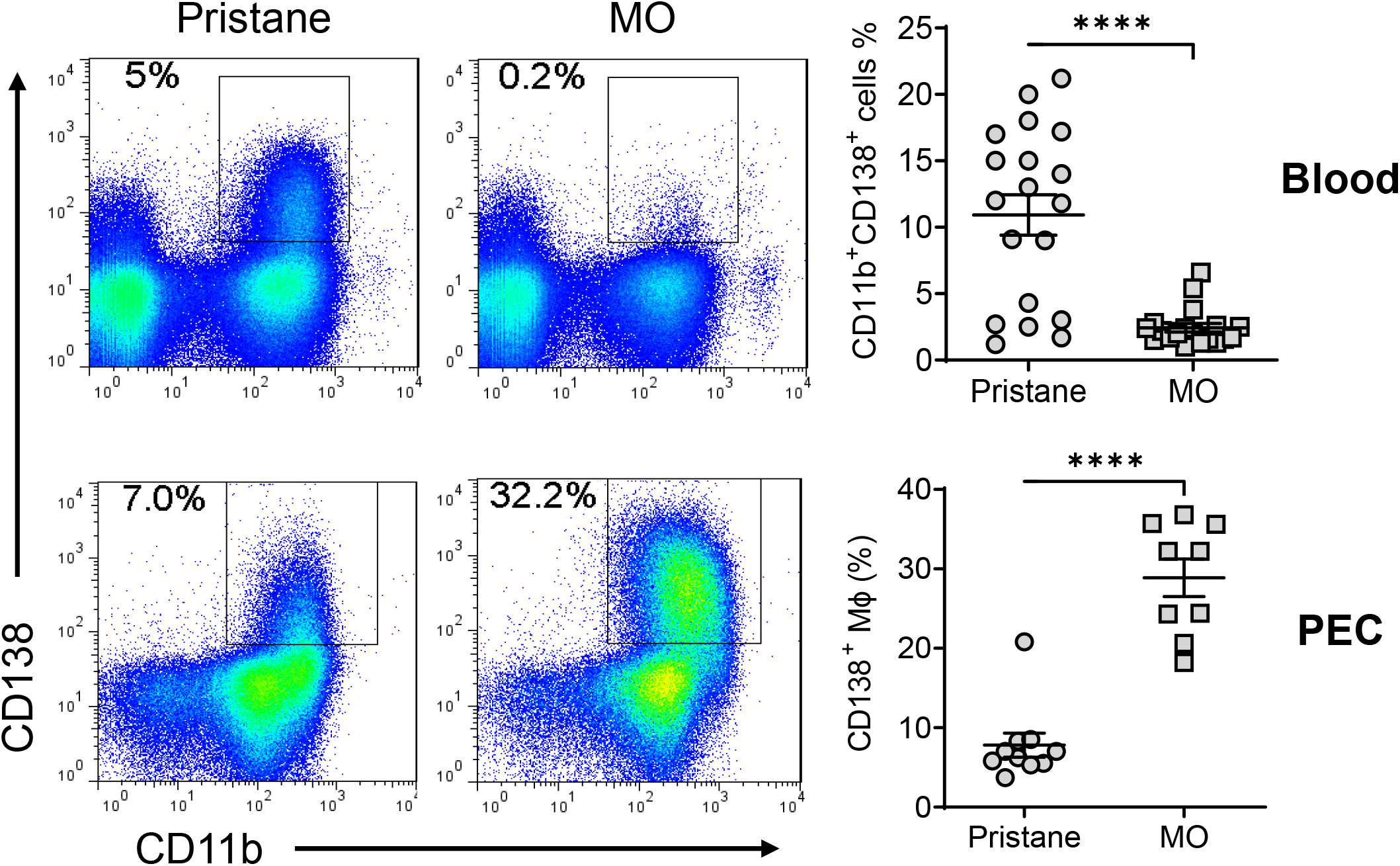
CD138^+^CD11b^+^ cells in the circulation and peritoneum. B6 mice were injected with either pristane or mineral oil (MO) and CD138 and CD11b surface expression on peripheral blood leukocytes and peritoneal exudate cells (PEC) was analyzed by flow cytometry at day-14. ****, P < 0.0001 (Student t-test).

Flow cytometry of CD11b^+^Ly6G^-^ cells from pristane-treated mice revealed three subsets in both the blood and the peritoneum **(Fig. 2A)**: CD11b^+^Ly6C^hi^CD138^-^ (R1); CD11b^+^Ly6C^lo/-^ CD138^-^ (R2); and CD11b^+^Ly6C^lo/-^CD138^+^ (R3). Ly6C staining was higher on the peritoneal R2/R3 cells vs. blood R2/R3 cells. In the blood of pristane-treated mice, all three subsets were CD43^+^ and TremL4^+^, consistent with a monocyte phenotype **(Fig. 2B-C)**. The blood R3 and R2 subsets expressed higher levels of CD43 and TremL4 than R1. R3 expressed higher levels than R2. CD115 was expressed at low levels, mainly on R3 cells **(Fig. 2D)**. As expected, CD62L, which is expressed by Ly6C^hi^ monocytes that have recently migrated out of the BM and not on Ly6C^lo^ monocytes (5), was expressed at considerably higher levels on R1 vs. R3 or R2 **(Fig. 2E)**. As classical monocytes are CD115^+^Ly6C^+^CD43^lo^ whereas non-classical monocytes are CD115^+^Ly6C^-^CD43^hi^TremL4^+^ (5), we concluded that the circulating R1 cells were classical (Ly6C^hi^) monocytes and the circulating R2 and R3 cells were subsets of non-classical monocytes. The circulating R2 subset was heterogeneous, with one population expressing high CD115, CD43, and TremL4 and a smaller population with low expression **(Fig. 2F)**. R1 also was heterogeneous, with two distinct populations expressing high and low levels of TremL4. Together the data suggest that peripheral blood of pristane-treated mice contains classical monocytes along with two subtypes of non-classical monocytes distinguished by the presence/absence of CD138 staining.

**Figure 2.**
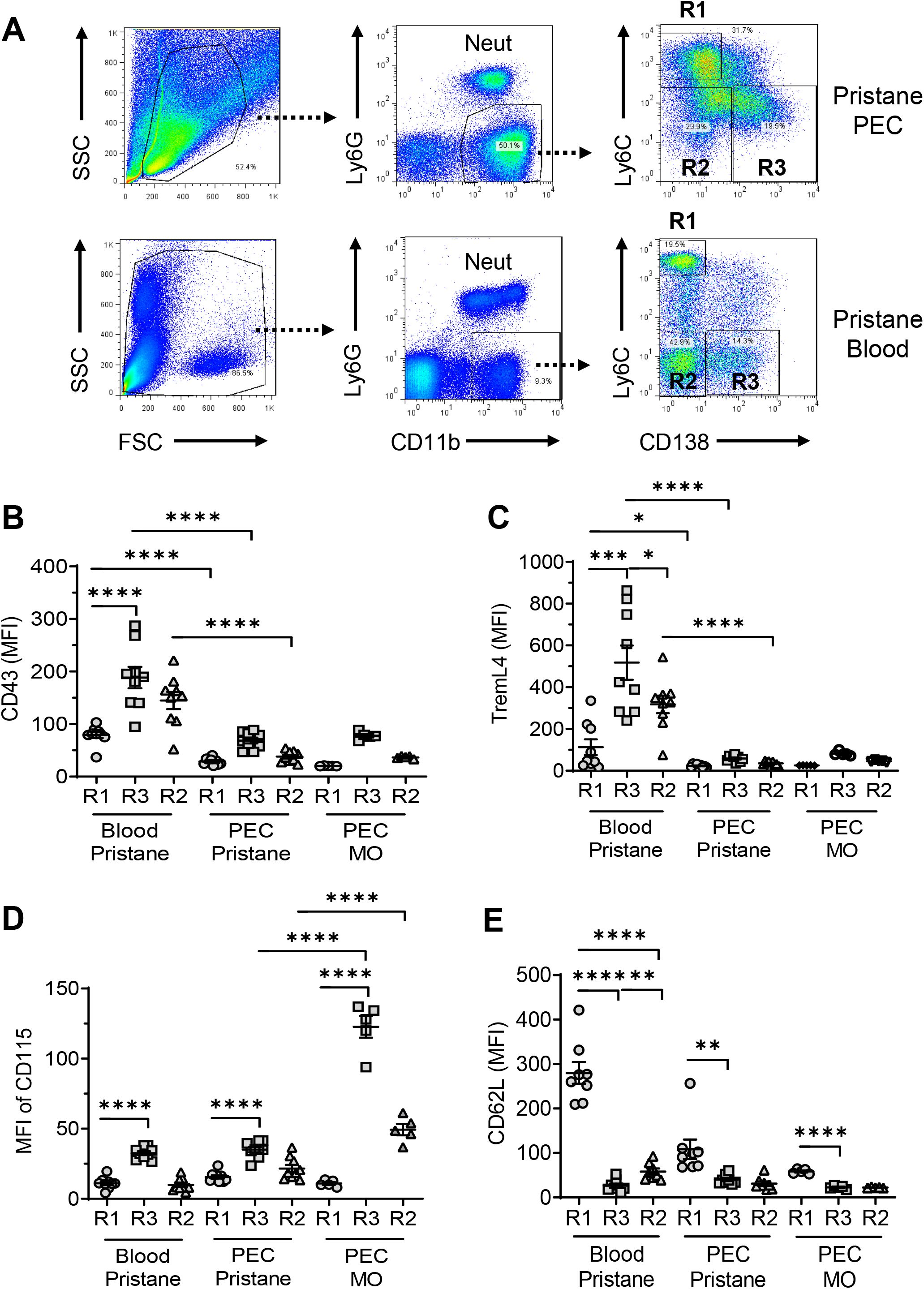
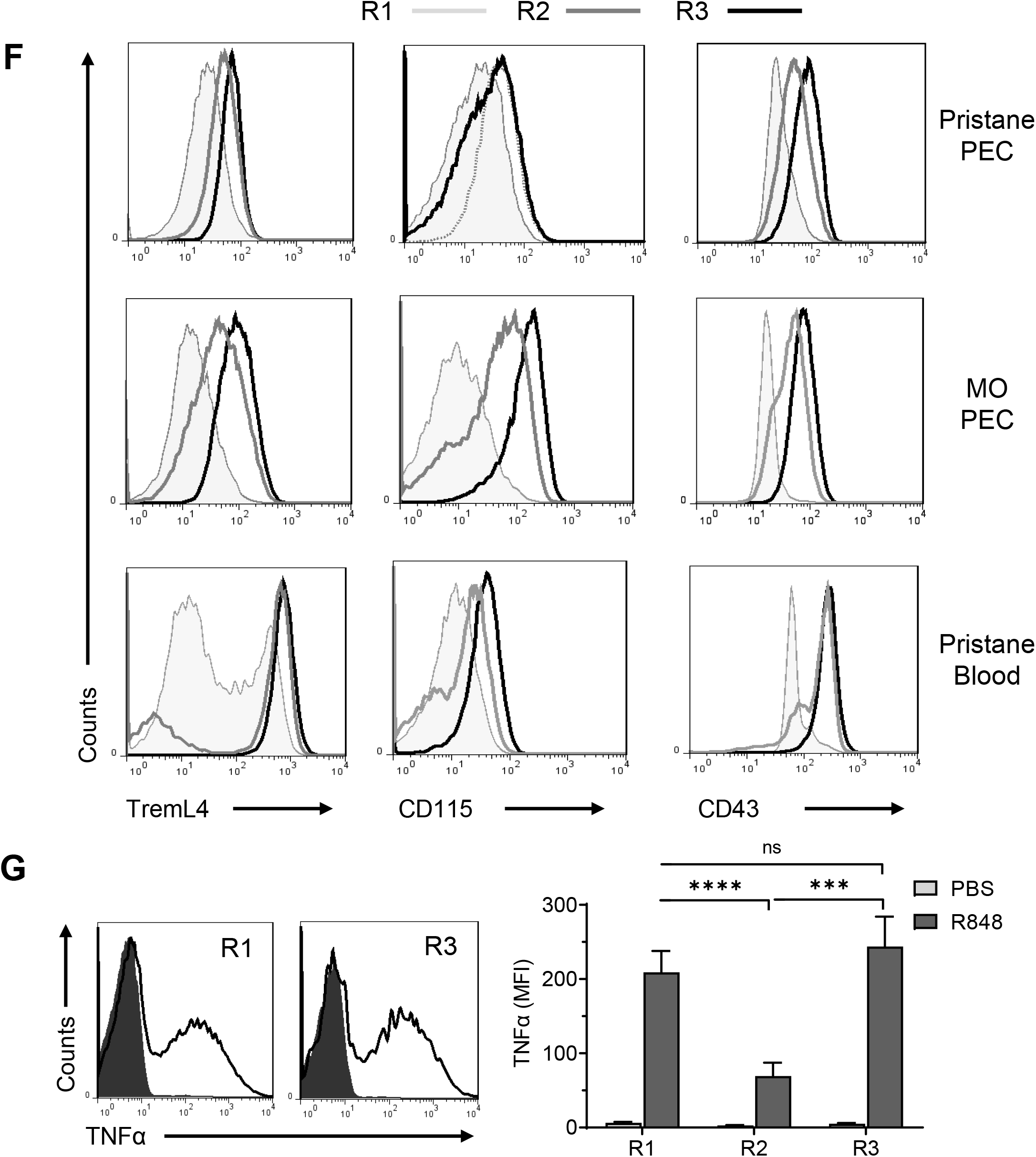
Flow cytometry of monocytes surface markers. Peripheral blood mononuclear cells and peritoneal exudate cells (PECs) from pristane- or MO-treated mice were stained with fluorescently labeled anti-CD11b, Ly6G, Ly6C, and CD138 antibodies. Mean fluorescence intensity (MFI) of CD43, TREML4, CD115, and CD62L was determined on cell subsets. **A,** Gating strategy for analysis of CD11b^+^Ly6G^-^ PEC and circulating mononuclear cells (blood). R1, CD11b^+^Ly6G^-^Ly6C^hi^CD138^-^ cells; R2, CD11b^+^Ly6G^-^Ly6C^-^CD138^-^ cells; R3, CD11b^+^Ly6G^-^Ly6C^-^CD138^+^ cells. **B-E,** staining intensity of CD43 (B), TREML4 (C), CD115 (D), and CD62L (E) in the R1, R2, and R3 subsets from peripheral blood or PEC from pristane or mineral oil (MO) treated mice. **F,** Representative histograms for TREML4, CD115, and CD43 staining. **G,** R848-stimulated TNFα production in R1, R2, and R3 monocytes from pristane-treated mice (14-d). Circulating leukocytes were incubated 5-h with R848 or PBS alone and then surface-stained with anti-CD11b, Ly6G, Ly6C, and CD138 antibodies and intracellularly stained with anti-TNFα. *Left,* histograms of intracellular TNFα staining in the R1 and R3 subsets. *Right,* TNFα staining of blood cells cultured 5-h with R848 or PBS. * P < 0.05; ** P < 0.01; *** P < 0.001; **** P < 0.0001 (Student t-test); ns, not significant.

Although non-classical monocytes are characteristically Ly6C^lo^, they have not been reported to express CD138. Since the Ly6C^lo^CD138^+^ subset expressed high levels of Treml4, a protein that amplifies TLR7-mediated signaling (15), we compared R848-stimulated TNFα production in CD138^+^ vs. CD138^-^ monocytes from pristane-treated mice **(Fig. 2G)**. As expected, Ly6C^hi^ monocytes (R1) exhibited strong intracellular TNFα staining 5-hr after culturing with R848. CD138^+^Ly6C^lo^ monocytes (R3) (which were Treml4^hi^) expressed similarly high levels of TNFα **(Fig. 2G)**. Both R1 and R3 expressed considerably more TNFα than CD138^-^Ly6C^lo^ (R2) monocytes.

### Peritoneal CD138^+^CD11b^+^ Mϕ

In contrast to the circulating R2 and R3 subsets, peritoneal R2 and R3 cells were strongly positive for the Mϕ markers F4/80 and CD64 as well as CD11b **(Figs. 3A-C)**. R2 and R3 cells from pristane-treated mice exhibited considerably stronger F4/80 and CD64 staining than R2 and R3 cells from MO-treated mice. Peritoneal R1 (Ly6C^hi^) cells in pristane-treated mice stained more weakly for F4/80, CD64, and CD11b than the R2 and R3 subsets **(Fig. 3A-C)**. But since these cells expressed little CD43, TremL4, or CD115 **(Fig. 2B-D)**, R1 also had a Mϕ-like phenotype. Thus, the peritoneal R1, R2, and R3 subsets all had a Mϕ-like phenotype and F4/80 and CD64 were higher in CD11b^+^CD138^+^Ly6C^-^ (R3) cells from pristane-vs. MO-treated mice.

**Figure 3.**
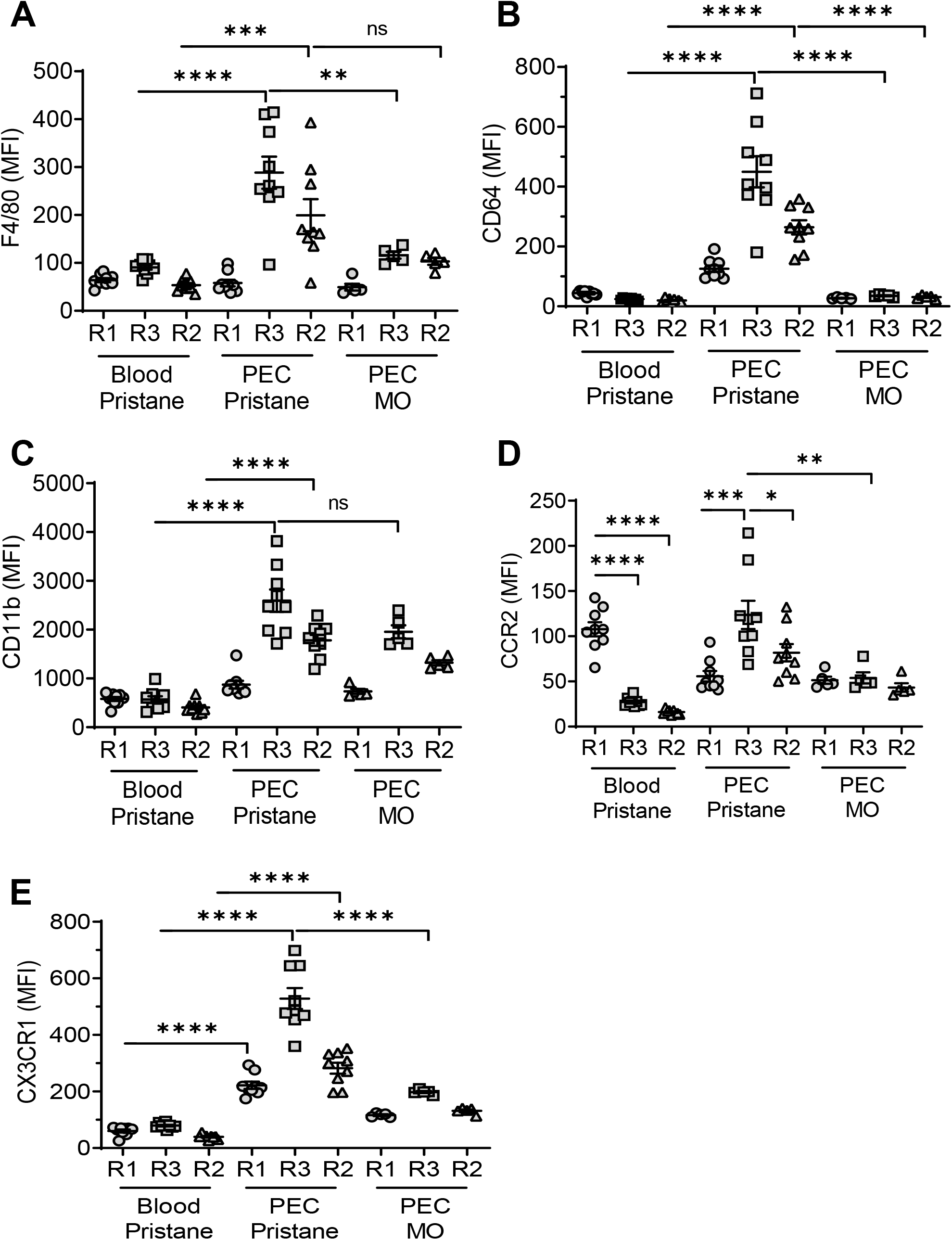
Flow cytometry of Mϕ markers in the blood and PECs. Peripheral blood mononuclear cells and PEC from pristane or MO treated mice were stained with fluorescently labeled anti-CD11b, Ly6G, Ly6C, and CD138 antibodies. CD11b^+^Ly6G^-^ myeloid subsets were gated as in Fig. 2A: R1, Ly6C^hi^CD138^-^, R2 Ly6C^lo^CD138^-^, R3 Ly6C^lo^CD138^+^. Mean fluorescence intensity (MFI) of F4/80 **(A)**, CD64 **(B)**, CD11b **(C)**, CCR2 **(D)**, and CX3CR1 **(E)**was determined. * P < 0.05; ** P < 0.01; *** P < 0.001; **** P < 0.0001 (Student t-test). ns, not significant.

### Circulating Ly6C^hi^ and CD138^+^ monocytes are Ccr2-dependent

Monocyte egress from the BM is defective in *Ccr2*-deficient mice (1) and Ccr2 is necessary for Ly6C^hi^ monocytes to migrate to the peritoneum in pristane-treated mice (6). After Ly6C^hi^ (R1) monocytes migrate to the peritoneum, they can down-regulate Ly6C. It is not known whether both Ly6C^lo^ peritoneal subsets (CD138^+^, R3 and CD138^-^, R2) are generated from Ly6C^hi^ precursors or if the peritoneal R3 subset is derived from circulating CD138^+^ monocytes. Circulating Ly6C^hi^ (R1) monocytes were strongly Ccr2^+^, whereas Ly6C^lo^CD138^-^ (R2) and Ly6C^lo^CD138^+^ (R3) monocytes expressed low levels **(Fig. 3D)**. In contrast, Ccr2 was expressed by all peritoneal subsets in pristane-treated mice, most strongly by R3. Ccr2 also was expressed by R1, R2, and R3 from MO-treated mice, but at lower levels **(Fig. 3D)**. Staining of circulating monocytes for Cx3cr1, which is expressed at high levels on Ly6C^lo^ monocytes but is involved in homing of both Ly6C^hi^ and Ly6C^lo^ monocytes (18, 19), was weak in pristane-treated mice, whereas peritoneal cells expressed higher levels, most prominently on R3 (CD138^+^ Mϕ) **(Fig. 3E)**. Expression was higher on peritoneal Mϕ from pristane- vs. MO-treated mice.

Circulating monocyte subsets varied with time after pristane treatment **(Fig. 4A)**. Ly6C^hi^ monocytes (R1) appeared first, about 5 days after pristane or MO treatment, while at the same time, the percentage of Ly6C^lo^CD138^-^ monocytes (R2) decreased. Circulating Ly6C^lo^CD138^+^ monocytes (R3) appeared in pristane-treated mice at around day-9 but only small numbers were seen in MO-treated mice **(Fig. 4A)**.

**Figure 4.**
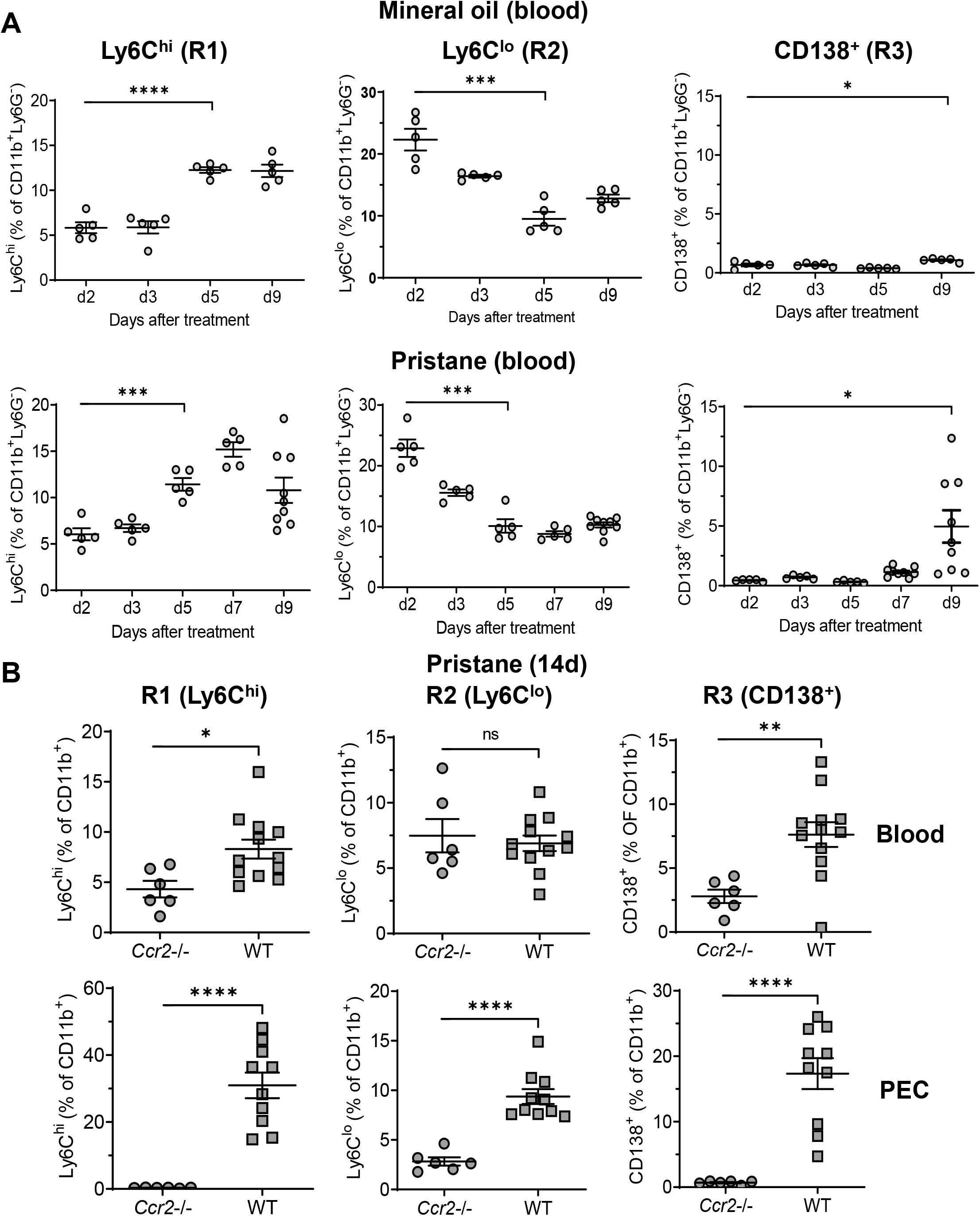
Ccr2 dependence of kinetics of monocyte egress from the BM. B6 mice were treated with either mineral oil or pristane and CD11b^+^Ly6G^-^ myeloid subsets in the blood and PEC were gated as in Fig. 2A. **A,**circulating Ly6C^hi^ (R1), Ly6C^lo^CD138^-^ (R2) and Ly6C^lo^CD138^+^ (R3) monocytes from mineral oil and pristane treated mice were assessed at 0-9 days after treatment by flow cytometry. Data are expressed as a percentage of CD11b^+^Ly6G^-^ cells. **B,**wild type B6 (WT) and B6 *Ccr2*-/- mice were treated with pristane. R1, R2, and R3 cells as a percentage of CD11b^+^Ly6G^-^ cells were measured in the blood and PEC by flow cytometry 14-d later. * P < 0.05; ** P < 0.01; *** P < 0.001; **** P < 0.0001 (Student t-test). ns, not significant.

In pristane-treated *Ccr2*-/- mice, circulating R1 and R3 cells were greatly reduced, but not absent **(Fig. 4B)**. In contrast, the percentages of circulating R2 cells were similar in *Ccr2*-/- vs. wild-type mice. In PECs, R1 and R3 cells were absent and R2 cells were present at lower levels in *Ccr2*-/- mice vs. wild-type **(Fig. 4B)**. Thus, the egress of R1 (Ly6C^hi^) and R3 (Ly6C^lo^CD138^+^) monocytes from the BM was Ccr2-dependent, whereas the circulating R2 monocyte subset (Ly6C^lo^CD138^-^) was unaffected by the absence of Ccr2. R1 and R3 monocytes appeared in the circulation with different kinetics. All three peritoneal Mϕ subsets were reduced in *Ccr2*-/- mice.

### CD138^+^ monocytes, but not CD138^+^ peritoneal Mϕ, are absent in Nr4a1-/- mice

Nr4a1 (Nur77) controls the differentiation of Ly6C^-^ monocytes (2) and regulates inflammatory responses in Mϕ (20). Peripheral blood cells and PECs from wild-type and *Nr4a1*-/- mice were analyzed by flow cytometry 14-d after pristane or MO treatment, gating on forward/side scatter and surface staining characteristics **(Fig. 5A)**. As expected (9), most peritoneal CD11b^+^Ly6G^-^ cells in MO-treated mice were Ly6C^lo^ and many were CD138^+^ **(Fig. 5A)**. In pristane-treated mice, the percentages of Ly6C^hi^ (R1), Ly6C^-^CD138^-^ (R2), and Ly6C^-^CD138^+^ (R3) peritoneal Mϕ were similar in *Nr4a1*-/- and wild-type mice **(Fig. 5B)**. In contrast, although the percentages of circulating R1 and R2 monocytes were similar, *Nr4a1*-/- mice had markedly fewer circulating CD138^+^ (R3) monocytes than wild-type mice **(Fig. 5C)**.

**Figure 5.**
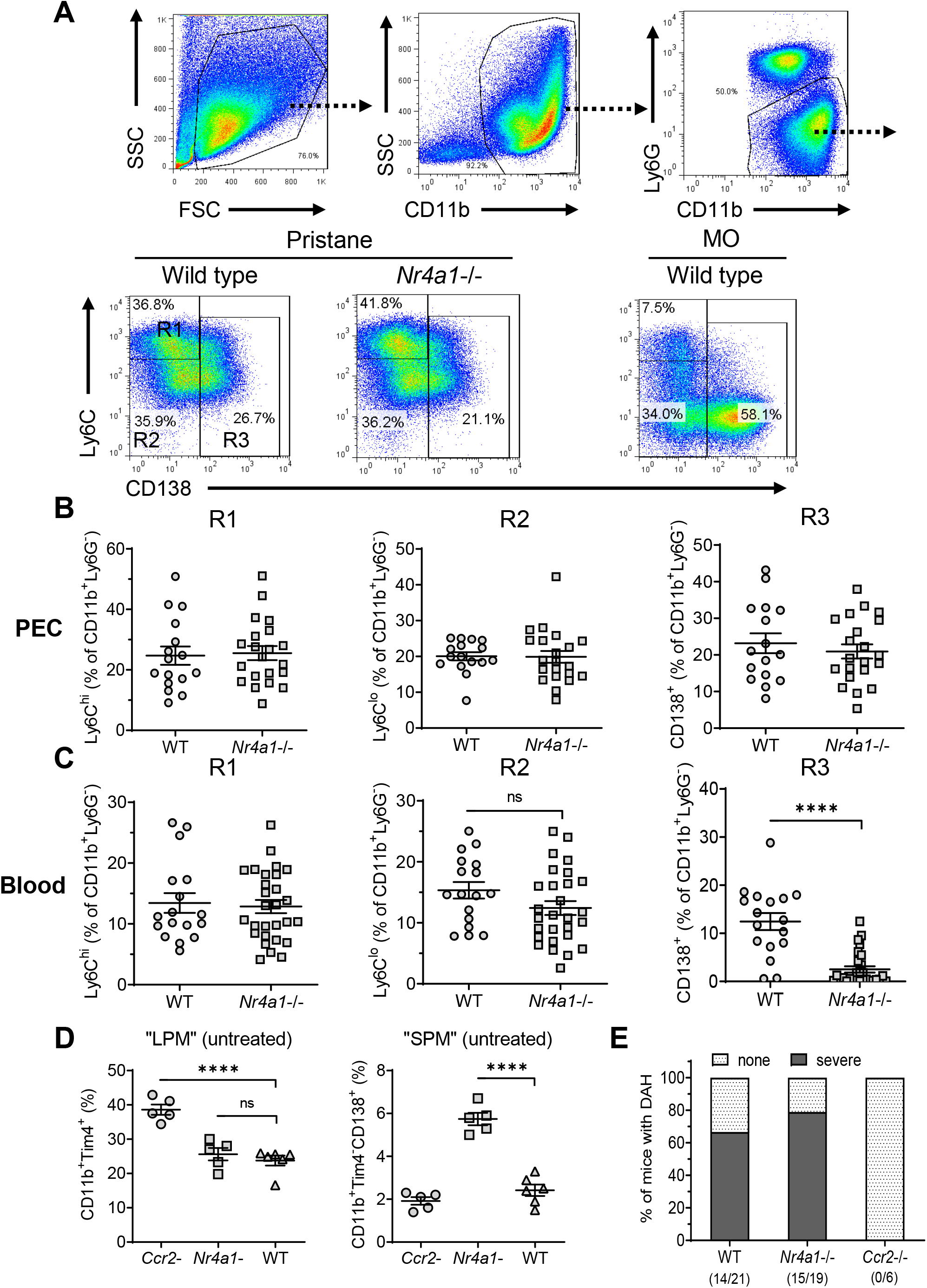
Monocyte/Mϕ subsets in Nr4a1-/- mice. **A,** Gating strategy. PECs from pristane- or MO-treated wild-type and *Nr4a1*-/- mice were stained with anti-CD11b, Ly6G, Ly6C, and CD138 and analyzed by flow cytometry. PECs were gated on the CD11b^+^Ly6G^-^ population. R1, Ly6C^hi^CD138^-^ PECs, R2, Ly6C^-^CD138^-^ PECs, R3, Ly6C^-^CD138^+^ PECs. Gating for circulating (blood) cells was similar. SSC, side scatter; FSC, forward scatter. **B,** R1, R2, and R3 subsets in the peritoneum (percentage of CD11b^+^Ly6G^-^ cells). **C,** R1, R2, and R3 subsets in the peripheral blood (percentage of CD11b^+^Ly6G^-^ cells). **D,** Flow cytometry of peritoneal cells in untreated *Ccr2*-/-, *Nr4a1*-/-, and wild-type (WT) mice. *Left,* CD11b^+^Tim4^+^ large peritoneal macrophages (LPM) as a % of total peritoneal cells. Right, CD11b^+^Tim4^-^CD138^+^ small peritoneal macrophages (SPM) as a % of total peritoneal cells. **E,** Frequency of diffuse alveolar hemorrhage (day-14) in pristane-treated WT, *Nr4a1*-/-, and *Ccr2*-/- mice. ****, P < 0.0001 (Student t-test). ns, not significant.

Along with CD11b^+^Tim4^+^ resident (yolk sac-derived) large peritoneal macrophages (LPM) (21), the uninflamed peritoneum contains CD11b^+^Tim4^-^CD138^+^ cells (22), which are probably “small peritoneal macrophages” (SPM) (21). The percentages of LPM in untreated wild-type vs. *Nr4a1*-/- mice were similar, whereas the percentage was higher in *Ccr2*-/- mice **(Fig. 5D)**, probably reflecting the absence of BM-derived cells. The CD138^+^ SPM subset was increased in *Nr4a1*-/- mice vs. wild-type and *Ccr2*-/- mice, suggesting that they originate from circulating BM-derived (Nr4a1-independent) Ly6C^+^ monocytes rather than Nr4a1-dependent CD138^+^ monocytes.

### Pristane-induced DAH is unaffected by absence of Nr4a1

Induction of DAH by pristane is abolished by eliminating monocytes and Mϕ with clodronate liposomes (8). Pristane-treated *Nr4a1*-/- mice developed DAH at a frequency comparable to wild-type controls **(Fig. 5E)**, suggesting that circulating CD138^+^ (R3) monocytes are dispensable for the induction of DAH. In contrast, DAH was abolished in *Ccr2*-/- mice suggesting that, as in the inflamed peritoneum, migration of Ly6C^hi^ monocytes from the BM to the lung is involved in the pathogenesis of DAH.

### Low TremL4 expression in Nr4a1-/- mice

TremL4 amplifies proinflammatory signaling through TLR7 by regulating the recruitment of MyD88 (14-17). It is expressed by Ly6C^lo^ monocytes and Nr4a1^+^Ly6C^hi^ monocytes committed to differentiate into Ly6C^lo^ monocytes (23). Circulating CD11b^+^Treml4^+^ monocytes increased in pristane-treated wild-type mice but were much less abundant in pristane-treated *Nr4a1*-/- mice vs. controls **(Fig. 6A)**. In both wild-type and *Nr4a1*-/- mice Treml4 staining intensity was highest in circulating CD138^+^ cells (R3) and was lower in CD138^-^Ly6C^lo/neg^ monocytes (R2A and R2B, respectively) **(Fig. 6B)**. Neutrophils were negative.

**Figure 6.**
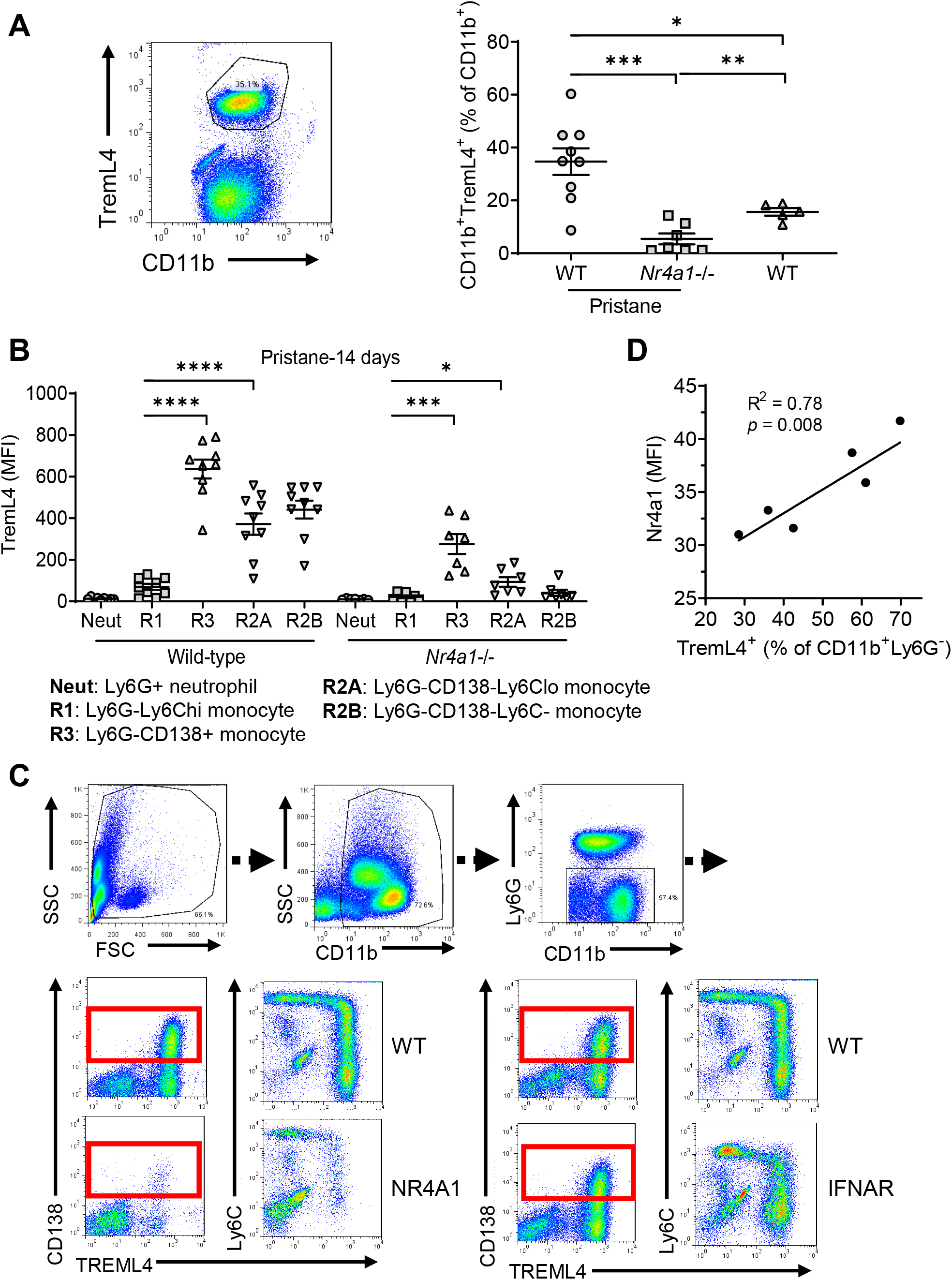
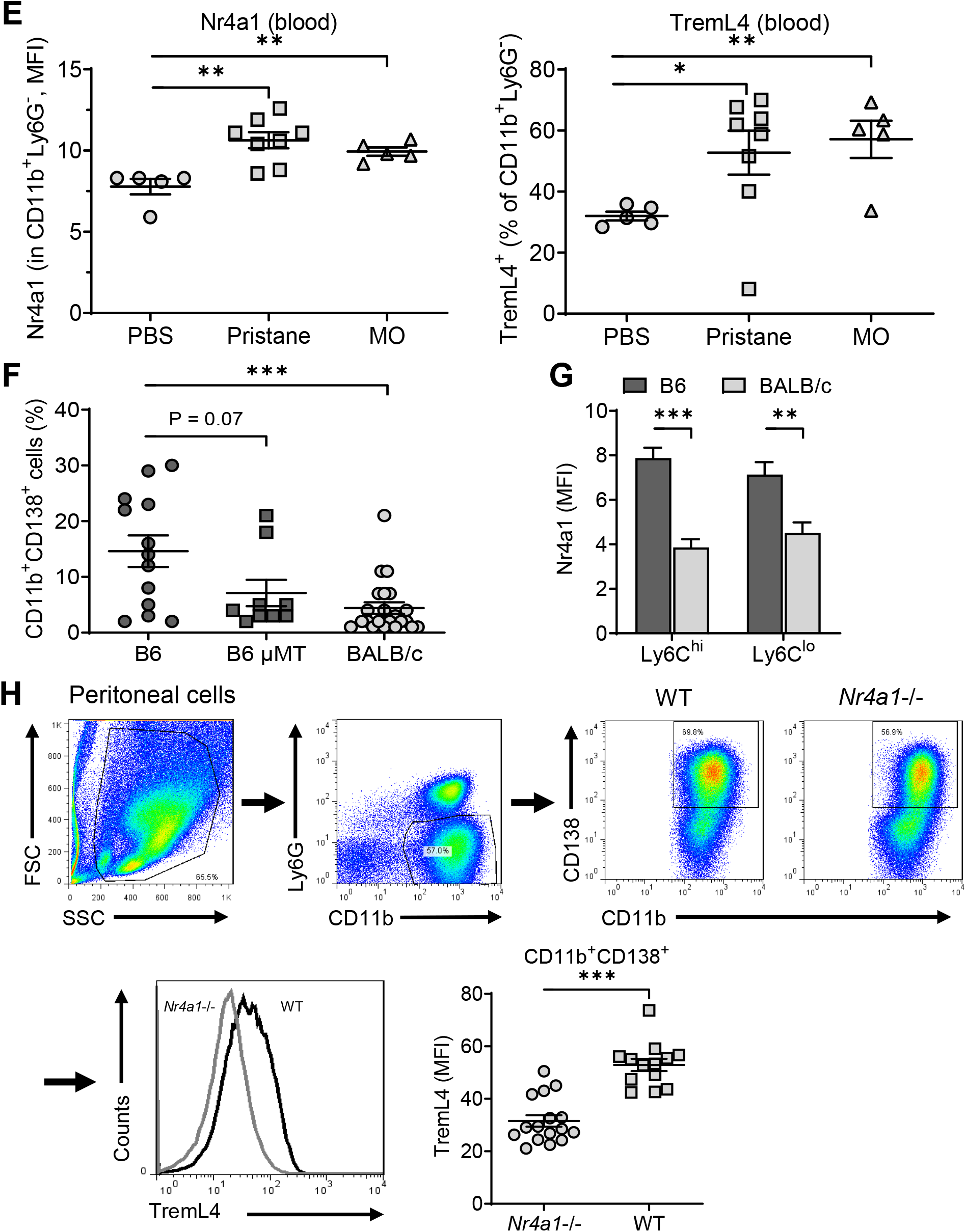
Low TremL4 expression in Nr4a1-/- mice. **A,** peripheral blood cells were stained with anti-CD11b and anti-Treml4 monoclonal antibodies. CD11b^+^Treml4^+^ cells as a percentage of total CD11b^+^ cells were determined by flow cytometry in pristane treated wild-type (WT) and *Nr4a1*-/- mice and in untreated WT mice. **B,** Wild-type and *Nr4a1*-/- mice were treated with pristane and CD11b^+^Ly6G^-^ myeloid subsets in the blood were gated as in Fig. 2A. CD11b^+^Ly6G^+^ neutrophils (percentage of total CD11b^+^ cells) and Ly6C^hi^ (R1), Ly6C^lo^ (R2), and CD138^+^ (R3) monocytes (percentage of CD11b^+^Ly6G^-^ cells) were determined by flow cytometry. **C,** *Left,* TremL4, CD138, and Ly6C staining of circulating CD11b^+^Ly6G^-^ cells from wild-type (WT) vs. *Nr4a1*-/- mice. *Right,* CD138, and Ly6C staining of circulating CD11b^+^Ly6G^-^ cells from wild-type (WT) vs. *Ifnar1*-/- mice. Nearly all CD138^+^ cells were TremL4^+^ (red boxes). **D,** Linear regression analysis of intracellular Nr4a1 and TremL4 staining in CD11b^+^Ly6G^-^ cells (R^2^ = 0.78, P = 0.008). **E,** Flow cytometry analysis of Nr4a1 (intracellular staining) and TremL4 (surface staining) in circulating (blood) CD11b^+^Ly6G^-^ cells from wild-type mice treated with PBS, pristane, or mineral oil (MO). **F,** Circulating CD138^+^ monocytes in blood from day-14 pristane-treated wild-type (B6), B cell deficient (B6 μMT), and BALB/c mice (flow cytometry). **G,** Intracellular Nr4a1 staining in the CD11b^+^Ly6C^hi^ and CD11b^+^Ly6C^lo^ circulating monocytes from pristane-treated B6 and BALB/c mice. **H,** TremL4 staining of peritoneal CD11b^+^CD138^+^ Mϕ from *Nr4a1*-/- mice and wild-type (WT) controls. * P < 0.05; ** P < 0.01; *** P < 0.001; **** P < 0.0001 (Student t-test).

Nearly all circulating CD138^+^ (R3) cells were TremL4^+^ **(Fig. 6C, boxes)**. Ly6C^hi^ (R1) cells may acquire Treml4 surface staining as they downregulate Ly6C and upregulate CD138 **(Fig. 6C, lower right)**. The acquisition of TremL4 and CD138 staining was attenuated in *Nr4a1*-/- mice. In contrast, although signaling through the type I interferon receptor (Ifnar1) blocks the downregulation of Ly6C in peritoneal Mϕ, *Ifnar1*-/- and wild-type mice had similar numbers of circulating CD138^+^TremL4^+^ monocytes **(Fig. 6C)**.

Intensity of Nr4a1 staining correlated with the percentage of circulating TremL4^+^CD11b^+^Ly6G^-^ cells **(Fig. 6D)**, suggesting that Nr4a1 might regulate TremL4. Consistent with that possibility, pristane and MO treatment both increased the fluorescence intensity of Nr4a1 as well as TremL4 in circulating CD11b^+^Ly6G^-^ cells **(Fig. 6E)**.

Since circulating Ly6C^lo^CD138^+^ monocytes were present in pristane-treated (develop DAH) but not MO-treated (do not develop DAH) B6 mice, we asked whether these Nr4a1-dependent cells are associated with susceptibility to pristane-induced DAH. Circulating Ly6C^lo^CD138^+^ monocytes were substantially lower in BALB/c mice (resistant to DAH induction) vs. B6 (DAH-susceptible) and there was a similar trend in pristane-treated B-cell-deficient (μMT) B6 mice, which also fail to develop DAH (8) **(Fig. 6F)**. Intracellular Nr4a1 staining (flow cytometry) was lower in circulating Ly6C^hi^ and Ly6C^lo^ monocytes from pristane-treated BALB/c vs. B6 mice **(Fig. 6G)**. Thus, high circulating, Nr4a1-dependent, Ly6C^lo^CD138^+^ monocytes and a high level of Nr4a1 protein in blood monocytes both were associated with the development of DAH.

Finally, we examined whether inflammation could increase TremL4 expression in PECs. Since there are relatively few CD138^+^Ly6C^lo^ (R3) Mϕ in pristane-treated mice, we examined PECs from MO-treated mice, gating on CD11b^+^Ly6G^-^CD138^+^ cells **(Fig. 6H)**. Although TremL4 staining was substantially lower on peritoneal CD138^+^ Mϕ vs. circulating CD138^+^ monocytes in wild-type mice **(R3, Fig. 2C)**, TremL4 staining of CD138^+^ peritoneal Mϕ was higher in wild-type vs. *Nr4a1*-/- mice **(Fig. 6H)**, suggesting that MO-induced peritoneal inflammation may induce TremL4 expression via Nr4a1.

### Pristane and TLR ligands induce TremL4 expression in RAW264.7 cells

TLR4 (LPS) and TLR7 (R848) ligands both activated the murine Mϕ cell line RAW264.7, as indicated by CD86 staining **(Fig. 7A)**and induced *Tnfa* mRNA expression **(Fig. 7B)**. R848 also stimulated Nr4a1 and TremL4 staining and there was a similar, but not statistically significant, trend in cells cultured with LPS **(Fig. 7A)**. *Nr4a1* mRNA expression increased rapidly, peaking ~1-hr after adding LPS or R848 **(Fig. 7B)**. *Treml4* and *Tnfa* mRNA peaked 3-hr after adding TLR ligands.

**Figure 7.**
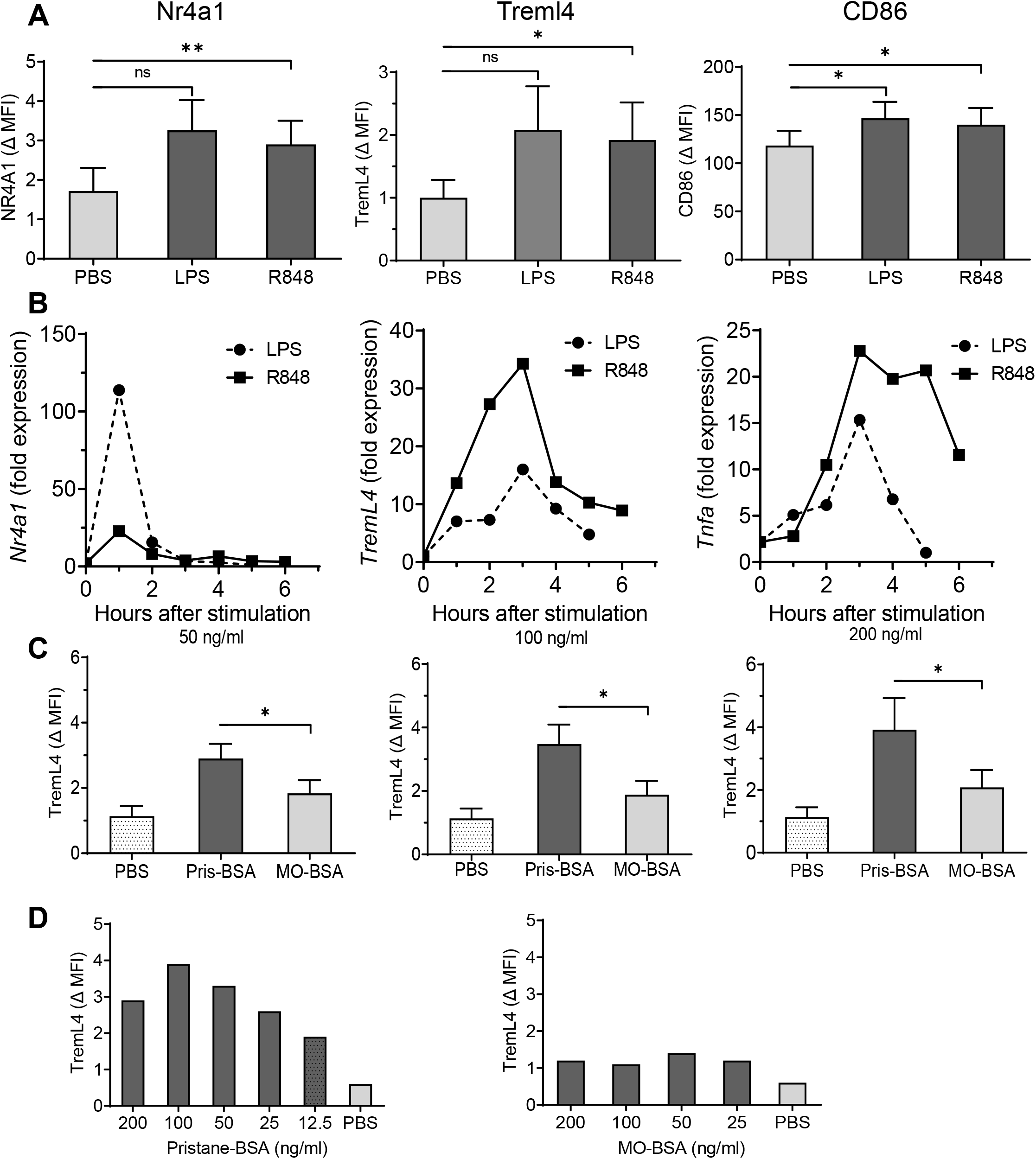
TremL4 and Nr4a1 protein expression in RAW264.7 Mϕ cell line. **A,** RAW264.7 cells were cultured for 24-hr with LPS (500 ng/mL), R848 (1 μg/mL), or PBS. Cells were then detached from the plastic wells and analyzed by flow cytometry for intracellular Nr4a1 and cell surface TremL4 and CD86 staining. **B,** time course of LPS- and R848-stimulated *Nr4a1*, *Treml4*, and *Tnfa* mRNA expression. **C,** RAW264.7 cells were cultured with pristane or mineral oil (MO) emulsified in bovine serum albumin (BSA) at doses of 50, 100, or 200 ng/ml for 24-hr or with PBS. Cells were detached from the wells and Treml4 expression was analyzed by flow cytometry. Data are expressed as change in mean fluorescence intensity (∆ MFI). **D,** dose-response for pristane-BSA (12.5-200 ng/ml) and MO-BSA (25-200 ng/ml) treatment expressed as ∆ MFI after 24-hr of culture.

TremL4 surface staining also was enhanced by culturing RAW264.7 cells with pristane (12.5-200 ng/ml emulsified in BSA) **(Fig. 7C-D)**. The effect was dose dependent and specific for pristane, as culture with MO did not enhance TremL4 staining.

### Nr4a1 regulates Treml4 in RAW264.7 cells and peritoneal CD138^+^ Mϕ

To further examine Nr4a1 regulation of *Treml4* expression, RAW264.7 cells were cultured with LPS in the presence of *Nr4a1* or control siRNA. As expected, *Nr4a1* siRNA downregulated LPS-stimulated *Nr4a1* mRNA expression at 1-hr **(Fig. 8A)**. *Nr4a1* siRNA also downregulated LPS-stimulated *Treml4* mRNA expression at 3-hr **(Fig. 8B)**.

**Figure 8.**
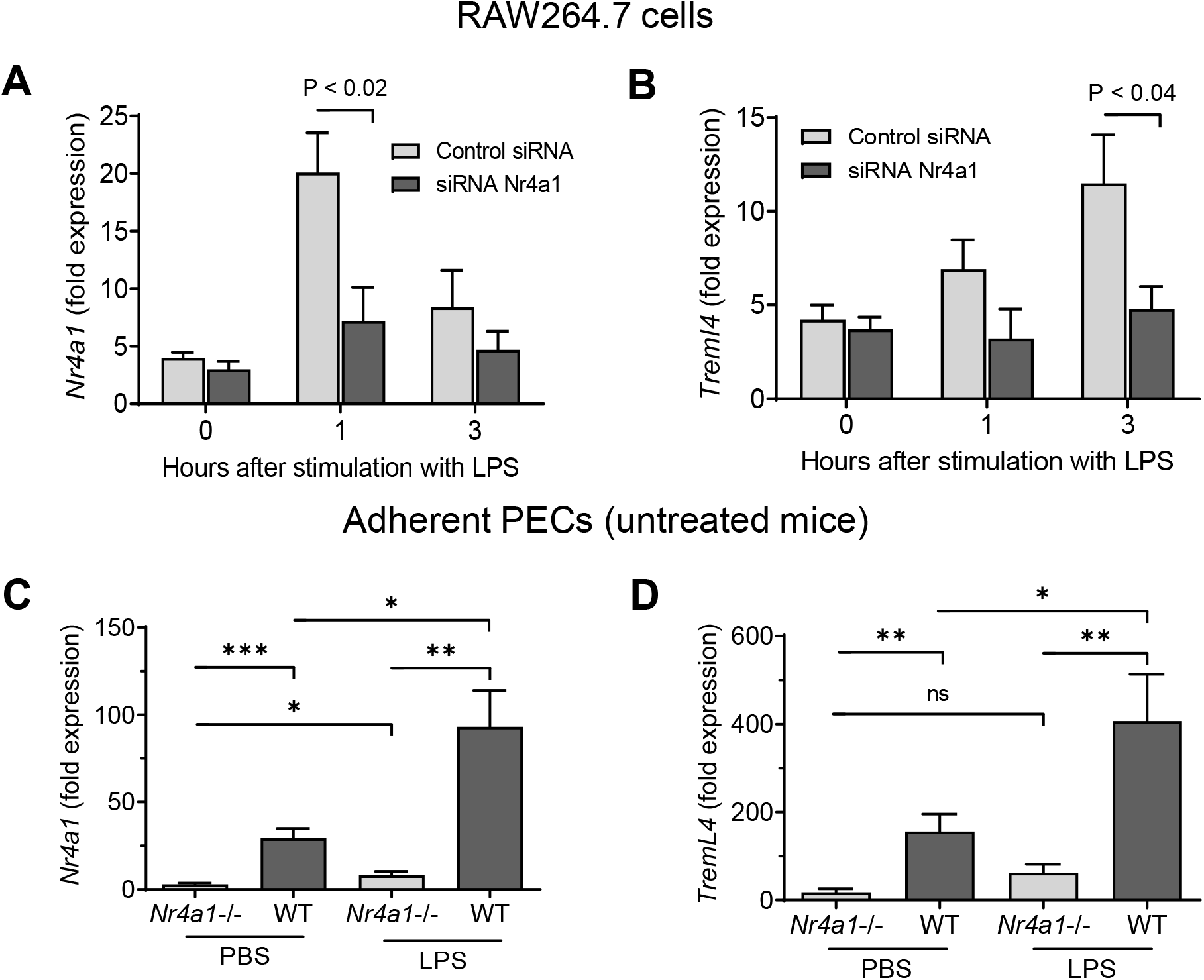
Nr4a1 regulates TremL4 mRNA and protein expression. **A-B,** RAW264.7 cells were transfected with 40 pmol *Nr4a1* or control siRNA and stimulated with LPS for 1-3 hours. Expression of *Nr4a1* **(A)** and *Treml4* **(B)** mRNA was quantified by qPCR at each timepoint. **C-D**, adherent peritoneal cells from untreated mice were cultured for 3-hours with LPS or PBS and then *Nr4a1* **(C)** and *Treml4* **(D)** mRNA was quantified by qPCR.

Peritoneal Mϕ from pristane-treated mice behaved similarly. TremL4 staining was lower in CD138^+^ Mϕ from *Nr4a1*-/- mice vs. wild-type controls **(Fig. 6H)**. As expected, *Nr4a1* expression was minimal in adherent peritoneal Mϕ from both PBS and LPS treated *Nr4a1*-/- mice compared with wild-type controls **(Fig. 8C)**. Unexpectedly, in *Nr4a1*-/- mice, *Nr4a1* increased slightly after LPS stimulation, probably due to the expression of a truncated fragment of the *Nr4a1* gene encoding part of the N-terminal domain in B6.129S2-*Nr4a1*^*tm1Jmi*^/J (*Nr4a1*-/-) mice (24). *Nr4a1* as well as *Treml4* expression was significantly higher in both PBS- and LPS-treated wild-type mice vs. *Nr4a1*-/- mice **(Fig. 8D)**.

## Discussion

Pristane-induced lupus in B6 mice is complicated by severe lung disease closely resembling DAH with antineutrophil cytoplasmic antibody (ANCA)-negative pulmonary vasculitis that is seen in SLE patients (8, 25). DAH and small vessel vasculitis are abolished by depleting monocytes/Mϕ with clodronate liposomes and are absent in mice lacking immunoglobulin or C3, suggesting that immune complexes are involved (8). Disease appears to result from pristane-induced lung microvascular injury (H Zhuang, submitted). Here, we examined the role of myeloid cell subsets in the pathogenesis of pristane-induced DAH. Lung disease was abolished in *Ccr2*-/- mice, suggesting that it depends on the influx of circulating BM-derived monocytes rather than activation of resident alveolar Mϕ. Circulating monocytes in pristane-treated mice included classical Ly6C^hi^ (inflammatory) monocytes and non-classical Ly6C^lo^ monocytes. Two subsets of Ly6C^lo^ monocytes were identified: CD138^-^ and CD138^+^. Ly6C^lo^CD138^+^ monocytes were dependent on the transcription factor Nr4a1, whereas the Ly6C^lo^CD138^-^ subset was Nr4a1-independent. Induction of DAH was associated with a wave of circulating Ly6C^lo^CD138^+^ monocytes 9-days after pristane treatment that was nearly absent in MO-treated B6 mice or pristane-treated BALB/c mice, neither of which develop DAH. However, *Nr4a1*-/- mice still developed DAH and vasculitis. In view of the important role of Nr4a1-dependent “patrolling” monocytes in monitoring vascular integrity (2), our data suggest that Ly6C^lo^CD138^+^ monocytes are generated in response to lung endothelial cell injury resulting from the influx of Ly6C^hi^Ccr2^+^ inflammatory monocytes.

### Two subsets of Ly6C^lo^ monocytes

Ly6C^lo^ (non-classical) monocytes develop from BM-derived Ly6C^hi^ precursors and are thought to be a single cell population (26, 27) that monitors the vascular endothelium and promotes the TLR7-dependent removal of damaged cells (11, 12). Development of this lineage requires the transcription factors Nr4a1 (2) and C/EBPβ (27). Circulating Ly6C^lo^ monocytes in pristane-treated mice consisted of CD138^-^ and CD138^+^ subpopulations, analogous to the peritoneal CD138^-^ and CD138^+^ Ly6C^lo^ Mϕ in these mice (22) **(Figs. 1–2)**. The two Ly6C^lo^ monocyte subsets differed functionally and developmentally. Like the *Nr4a1*-dependent Ly6C^lo^ monocytes reported previously (2), CD138^+^ monocytes were deficient in *Nr4a1*-/- mice **(Fig. 5C)**. In contrast, although strongly positive for monocyte markers CD43 and Treml4 **(Fig. 2)**, Ly6C^lo^CD138^-^ monocytes were Nr4a1-independent, and are a distinct subset. Nr4a1-independent Ly6C^lo^ monocytes have been encountered before but their function is unclear (11). Consistent with a BM-derived Ly6C^hi^ precursor (3, 26–28), circulating Ly6C^lo^CD138^+^ monocytes were absent in pristane-treated *Ccr2*-/- mice **(Fig. 4B)**.

Although Ly6C^lo^CD138^-^ monocytes circulated with similar kinetics in pristane- and MO-treated mice, Ly6C^lo^CD138^+^ monocytes only appeared 9-d after pristane treatment and were nearly absent in MO-treated mice **(Fig. 4)**. The Ly6C^lo^CD138^+^ subset might promote persistence rather than resolution of inflammation, as suggested by the higher TNFα production of Ly6C^lo^CD138^+^ vs. Ly6C^lo^CD138^-^ monocytes **(Fig. 2)**. Alternatively, since Nr4a1-dependent Ly6C^lo^ monocytes monitor endothelial cell damage (11), the Ly6C^lo^CD138^+^ subset could be generated in response to microvascular injury. Nr4a1-dependent Ly6C^lo^ monocytes promote TLR7- and neutrophil-dependent necrosis and removal of damaged endothelial cells. Analogous to the neutrophil “left-shift” seen in infections (29), circulating Ly6C^lo^CD138^+^ monocytes might be a clinically-relevant indicator of endothelial damage. Consistent with the possibility that the number of these cells reflects the severity of endothelial damage, pristane causes more severe lung microvascular injury in B6 vs. BALB/c mice (H Zhuang, et al. Submitted) and circulating Ly6C^lo^CD138^+^ monocytes were increased pristane-treated B6 vs. BALB/c mice **(Fig. 6F)**.

### What is the role of CD138?

The function of CD138 (syndecan-1) in myeloid cells is uncertain. Although CD138 was expressed on a subset of Ly6C^lo^ monocytes and Mϕ, *Sdc1* is not among the genes differentially expressed in Ly6C^hi^ vs. Ly6C^lo^ monocytes from non-pristane-treated-mice (27). There are several plausible explanations. First, the anti-CD138 monoclonal antibody we used might cross-react with another target in myeloid cells. However, clone 281-2 is a standard antibody for flow cytometry of mouse CD138 that recognizes the extracellular domain of CD138 on plasma cells, epithelial cells, and early B cells (30). Although transcriptional profiling of Ly6C^lo^CD138^+^ monocytes was not done, flow-sorted CD138^+^ Mϕ express high levels of *Sdc1* mRNA (31) and other investigators also have reported CD138 mRNA and protein expression in peritoneal Mϕ (32). Alternatively, CD138 might be regulated post-transcriptionally in monocytes, as in peritoneal Mϕ (32) and B cells (33). However, CD138 protein expression correlates with the abundance of *Sdc1* transcripts in peritoneal Mϕ [(31); S Han, not shown]. A third possibility is that *Sdc1* is not normally expressed in monocytes, but is induced by pristane in response to endothelial injury (H Zhuang, et al. Submitted). It may be worthwhile to see if circulating CD138^+^ monocytes are present in other models of vascular injury.

CD138 is a heparan sulfate-containing membrane proteoglycan that binds extracellular matrix (ECM) proteins and promotes integrin activation, wound healing, cell adhesion/migration, endocytosis, and fibrosis (34). Its functions in normal and neoplastic epithelial, endothelial, and stromal cells have been studied extensively, but little is known about its role in myeloid cells. The extracellular domain binds type IV collagen and laminins (34–37), which are components of the basement membrane separating the alveolar capillary endothelium and the epithelium (38). In chronic inflammation, monocytes attach to the vascular endothelium, migrate to the subendothelial space, and reversibly attach to basement membrane laminin via integrins (39, 40). It will be of interest to investigate whether expression of CD138 by Ly6C^lo^ monocytes promotes migration and/or attachment to the basement membranes of damaged blood vessels.

### CD138^+^ monocytes are not precursors of peritoneal CD138^+^ Mϕ

Migration of Ly6C^hi^ monocytes to the peritoneum in pristane-treated mice requires Ccr2 (6). Consistent with a previous report (11), circulating Ly6C^hi^ monocytes were reduced by about 50% in pristane-treated *Ccr2*-/- mice **(Fig. 4B)**. However, in the peritoneum, the absence of *Ccr2* eliminated both Ly6C^hi^ and Ly6C^lo^CD138^+^ Mϕ **(Fig. 4B)**. The inability of the remaining circulating Ly6C^hi^ monocytes to enter the peritoneum suggests that both BM egress and migration to the peritoneum are Ccr2-dependent. Following monocyte/Mϕ depletion with clodronate liposomes, the peritoneum is re-populated by Ly6C^hi^ monocytes, which can differentiate into Ly6C^lo^ Mϕ (6). The absence of both Ly6C^hi^ and CD138^+^ Mϕ in the peritoneum of *Ccr2*-/- mice suggests that CD138^+^ Mϕ are derived from Ly6C^hi^ monocyte precursors.

### Treml4 is transcriptionally regulated by Nr4a1

*Nr4a1* (Nur77) is a member of the nuclear receptor 4A subfamily that regulates monocyte and Mϕ differentiation (41). Nr4a1 is highly expressed in Ly6C^lo^ monocytes and *Nr4a1*-/- mice lack circulating Ly6C^lo^ monocytes (2, 42). Our data suggest that *Treml4* is one of the genes regulated by Nr4a1. *Nr4a1* correlated with *Treml4* mRNA and protein in pristane-treated mice and in TLR ligand- or pristane-treated RAW264.7 cells **(Fig. 6–8)**. Conversely, *Nr4a1* knockdown decreased *Treml4* expression in RAW264.7 cells and *Treml4* expression was lower in adherent Mϕ from *Nr4a1*-/- mice vs. controls **(Fig. 8)**. Although Nr4a1 appears to transcriptionally regulate *Treml4*, additional factors are likely to be involved, in light of the expression of low levels of *Treml4* in Ly6C^lo^CD138^+^ and Ly6C^lo^CD138^-^ monocytes and Mϕ from *Nr4a1*-/- mice **(Fig. 6)**.

Relevant to the role of Ly6C^lo^ patrolling monocytes in monitoring for endothelial damage, TremL4 binds late apoptotic and necrotic cells and sensitizes cells to TLR7 signaling by recruiting MyD88 to endosomes (14, 15). *Treml4*-/- MRL/*lpr* mice have lower autoantibody and interferon-α production and improved survival vs. *Treml4*+/+ controls (15), suggesting that Treml4 plays a causal role in lupus nephritis by upregulating TLR7-driven interferon production. There also is evidence that patrolling monocytes mediate glomerular inflammation in murine lupus nephritis (43).

Genetic polymorphisms that increase *TREML4* mRNA expression in human peripheral blood cells are associated with the progression and extent of coronary atherosclerotic lesions (44). However, a causal relationship has not been established and most evidence suggests that patrolling monocytes are protective in atherosclerosis (20, 45). It is possible that upregulation of TLR7 signaling by Treml4 has a dual effect, on one hand promoting vascular integrity by enhancing the removal of damaged endothelial cells, and on the other enhancing vascular inflammation if the endothelial cell damage cannot be resolved.

In summary, we identified two subsets of Ly6C^lo^ non-classical monocytes in pristane-treated mice, one CD138^+^ and Nr4a1-dependent and the other CD138^-^ and Nr4a1-independent. Although CD138 expression has not been reported previously on Ly6C^lo^ monocytes, the Nr4a1 dependence of Ly6C^lo^CD138^+^ cells strongly suggests they are patrolling monocytes involved in monitoring vascular integrity. We conclude that pristane-induced lung microvascular lung injury stimulates the production of these cells in an ineffectual effort to maintain vascular integrity in the face of ongoing endothelial damage. It remains to be determined whether Ly6C^lo^CD138^+^ monocytes can be detrimental when vascular damage is unresolved.

## Materials and Methods

### Mice

C57BL/6 (B6), B6.129S4-*Ccr2*^*tm1Ifc*^/J (*Ccr2*-/-), B6.129S2-*Nr4a1*^*tm1Jmi*^/J (*Nr4a1*-/-), and B6(Cg)-Ifnar1^tm1.2Ees/J^ (*Ifnar1*-/-) mice (Jackson Laboratory, Bar Harbor, ME) maintained under specific pathogen free conditions were injected i.p. with 0.5 ml of pristane (Sigma-Aldrich, St. Louis, MO) or mineral oil (MO, E.R. Squibb & Sons, New Brunswick, NJ) as described (8). There were 5-15 mice female mice age 8-12 weeks per group unless otherwise noted. Experiments were repeated at least twice. Peritoneal exudate cells (PECs) were collected by lavage 14-days later and heparinized blood was collected from the heart 3-14 days after pristane injection. DAH was assessed as described (8). This study followed the recommendations of the Animal Welfare Act and US Government Principles for the Utilization and Care of Vertebrate Animals and was approved by the UF IACUC.

### Flow cytometry

Flow cytometry of peritoneal cells was performed using anti-mouse CD16/32 (Fc Block, BD Biosciences, San Jose, CA) before staining with primary antibody or isotype controls. Peripheral blood (100 μL) was incubated 30-min in the dark with monoclonal antibodies specific for surface markers. After surface-staining, cells were fixed/permeabilized for 5-min (Fix-Perm buffer, eBioscience, San Diego, CA), washed and stained intracellularly for 20-min with anti-Nr4a1 monoclonal antibodies. The cells were washed and re-suspended in PBS for flow cytometry. Monoclonal antibodies are listed in **Table S1**.

### R848 stimulation of murine monocytes

Blood was collected by cardiac puncture from B6 mice 14-d after pristane treatment. Erythrocytes were lysed with erythrocyte lysis buffer (Qiagen) and washed with PBS. Leukocytes were re-suspended in complete DMEM + 10% fetal bovine serum in the presence or absence of R848 (1 μg/mL, Sigma-Aldrich) and cultured in a 5% CO2 atmosphere at 37°C for 5-hours. The cells were prepared for flow cytometry as above using anti-Ly6C-FITC, Ly6G-APC-Cy7, CD138-PE, and CD11b-BV-421 (surface staining) followed by intracellular staining with anti-TNFα-APC antibodies. Ly6G^+^ cells were gated out and intracellular TNFα staining was measured in CD11b^+^Ly6C^hi^CD138^-^, CD11b^+^Ly6C^lo^CD138^-^, and CD11b^+^Ly6C^hi^CD138^+^ cells (R1, R2, and R3, respectively).

### RAW264.7 cell culture with TLR ligands

RAW264.7 cells (murine macrophage cell line, ATCC, Manassas, VA) were seeded in 24 well non-attachment plates (5 × 10^5^ cells/well) and cultured for 24-h in complete DMEM + 10% fetal bovine serum with or without LPS (1 μg/mL) or R848 (1 μg/mL). The cells were washed with PBS and analyzed by flow cytometry. Cells were surface stained for TremL4 and CD86 and intracellularly stained for Nr4a1. For qPCR analysis, RAW264.7 cells were seeded in 6-well plates (3 × 10^5^ cells/well) and cultured overnight in complete DMEM + 10% FBS. On the following day, the cells were stimulated with LPS (1 μg/mL) or R848 (1 μg/mL) or PBS for 1, 2, 3, 4, 5, or 6 hours. Total RNA was obtained using the QIAamp RNA blood Mini Kit (Qiagen, Germantown, MD), and cDNA was synthesized using the Superscript II First-Strand Synthesis kit (Invitrogen, Carlsbad, CA). SYBR Green qPCR analysis was performed using the CFX Connect Real-Time system (Bio-Rad, Hercules, CA). Gene expression was normalized to 18S RNA and the expression level was calculated using the 2^−ΔΔCt^ method.

Primer sequences were as follows: *Nr4a1* forward: AGCTTGGGTGTTGATGTTCC; *Nr4a1* reverse: ATGCGATTCTGCAGCTCTT; *Treml4* forward: CTGGAGGTACTCACAACTGCT; *Treml4* reverse: GGCTCTGTCCTACCATTCTATGA; *Tnfa* forward: AGGAGGAGTCTGCGAAGAAGA; *Tnfa* reverse: GGCAGTGGACCATCTAACTCG; 18S rRNA forward: TGCCATCACTGCCATTAAGG; reverse: TGCTTTCCTCAACACCACATG.

### RAW264.7 cell culture with pristane

Pristane or mineral oil (1 ml) was added to PBS (9 ml) containing 100 mg/ml BSA in a 15-ml polypropylene tube and rotated for 48-h at 4°C. The surface layer of unincorporated hydrocarbon oil was aspirated at the end of the incubation. The amount of pristane incorporated using this method was calculated as described (8) and adjusted to approximately 1 μg/ml. RAW264.7 cells were seeded in 24-well non-attachment plates (3 × 10^5^ cells/well) and cultured for 24-h in complete DMEM + 10% FBS with/without hydrocarbon oils at concentrations ranging from 12.5-200 ng/ml. Then cells were washed with PBS and analyzed by flow cytometry for surface TremL4 and intracellular Nr4a1.

### Nr4a1 gene silencing

RAW264.7 cells were seeded in 6-well plates (2 × 10^5^ cells/well) and cultured overnight in antibiotic-free complete DMEM + 10% FBS. On the following day the cells were transfected with Nr4a1 (Nur77) siRNA (sc-36110 from Santa Cruz Biotechnology) or control siRNA-A (sc-37007, Santa Cruz Biotechnology, Dallas RX). siRNA (40 pmol from a 10 μM stock in sterile water) was added to 100 μl of Opti-MEM medium and transfected using siRNA transfection reagent (sc-29528, Santa Cruz Biotechnology) following the manufacturer’s instructions. Transfected cells were incubated at 37°C for 48-h, and then stimulated with LPS (1 μg/ml) for 1, 3 or 6-h. Total RNA was collected from the cells using the QIAamp RNA blood Mini Kit (Qiagen) and cDNA was synthesized using the Superscript II First-Strand Synthesis kit (Invitrogen). SYBR Green qPCR analysis was performed using the CFX Connect Real-Time system (Bio-Rad). Gene expression was normalized to 18S RNA and the expression level was calculated using the 2^-^ΔΔCt method. Primer sequences were as above.

### Peritoneal cell culture with TLR ligands

PECs were harvested from untreated wild-type B6 and *Nr4a1*-/- mice and allowed to adhere to plastic wells (6-well plates, 2 × 10^6^ cells/well) for 1 hour in AIM V serum free medium (ThermoFisher Scientific, Waltham, MA). Non-adherent cells were washed off with PBS and the adherent cells were cultured for 3-hrs in complete DMEM + 10% FBS medium containing LPS (1 μg/ml in PBS) or PBS alone. *Nr4a1* and *Treml4* expression was quantified by qPCR as above.

### Statistical analysis

Statistical analyses were performed using Prism 6.0 (GraphPad Software, San Diego, CA). Differences between groups were analyzed by two-sided unpaired Student *t* test unless otherwise indicated. Data were expressed as mean ± SD. *p* < 0.05 was considered significant. Experiments were repeated at least twice.

## Acknowledgements

This work was supported by the National Institutes of Health (NIAMS) grant number R01-AR44731 (WR) and by Department of Medicine (HZ).

---

**Table S1.** Monoclonal antibodies.

| **Specificity (clone)** | **Fluorochrome\*** | **Source** |
| --- | --- | --- |
| Mouse CD138 (281-2) | PE; APC; APC-Cy7 | BioLegend |
| Mouse CD11b (M1/70) | BV421, Pacific Blue | BioLegend |
| Mouse Ly6C (HK1.4) | APC-Cy7; Alexa Fluor 488 | BioLegend |
| Mouse Ly6G (1A8) | APC-Cy7; PE | BioLegend |
| Mouse CD43 | FITC | Biolegend |
| Mouse TREML4 | PE | Biolegend |
| Mouse CD115 | APC-Cy7 | Biolegend |
| Mouse F4/80 | PE, BV421 | Biolegend |
| Mouse CD62L | PE | BD pharmingen |
| Mouse CD64 | PE | BD pharmingen |
| Mouse CCR2 | FITC | Biolegend |
| Mouse Tim4 | APC, PE | Biolegend |
| Mouse CX3CR1 (SA011F11) | FITC | BioLegend |
| Mouse NR4A1 | PE | Miltenyi |
| Mouse IFNAR1 (MAR1-5A3) | APC | BioLegend |
| Mouse CD86 | APC-Cy7 | Biolegend |
| Mouse TNFα | APC, PE | Biolegend |

\* PE, phycoerythrin; APC, allophycocyanin; FITC, fluorescein isothiocyanate; BV, brilliant violet; APC-Cy7, APC/cyanine 7.

